# GbyE: A New Genome Wide Association and Prediction Model based on Genetic by Environmental Interaction

**DOI:** 10.1101/2023.05.17.541129

**Authors:** Xinrui Liu, Mingxiu Wang, Jie Qin, Yaxin Liu, Jincheng Zhong, Jiabo Wang

## Abstract

Nowadays, studies on genetic by environment interactions (G×E) are receiving increasing attention because of its theoretical and practical importance in explaining individual behavioral traits. Use information from different environments to improve the statistical power of genome-wide association and prediction in the hope of obtaining individuals with better breeding value is the most expedient way. However, there are significant challenges when performing genome-wide association studies (GWAS) and genomic selection (GS) using multiple environments or traits, mainly because most diseases and quantitative traits have numerous associated loci with minimal effects. Therefore, this study constructed a new genotype design model program (GbyE) for genome-wide association and prediction using Kronecker product, which can enhance the statistical power of GWAS and GS by utilizing the interaction effects of multiple environments or traits. The data of 282 maize, 354 yaks and 255 peaches were used to evaluate the power of the model at different levels of heritability and genetic correlation. The results show that GbyE can provide higher statistical power for the traditional GWAS and GS models in any heritability and genetic correlation, and can detect more real loci. In addition, GbyE has increased statistical power to three Bayesian models (BRR, BayesA, and BayesCpi). GbyE can make full use of multiple environment or trait informations to increase the statistical power of the model, which can help us understand the G×E and provide a method for predicting association loci for complex traits.

## 1. Introduction

Genetic by environment interactions (G×E) are of great theoretical and practical importance in explaining individual behavioral traits, which has led to the fact that G×E studies are gaining more and more attention. G×E can be understood as the influence of genetic factors on the susceptibility to environmental factors. In-depth study of G×E contributes to a deeper understanding of the relationship between individual growth or living environment and phenotypes. Almost all human diseases are currently studied to be associated with genetic factors at the molecular or cellular level, and non-genetic factors (environmental factors) may also be important contributors to human diseases. Of course, the goal of most researchers exploring the inevitable interactions between organisms and the environment is to understand the mechanisms behind complex diseases as well as complex quantitative traits. In the study of human diseases, whether they are common, complex or rare, we mostly consider them to be the result of a combination of genes, environmental factors and their interactions. By understanding the joint analysis between genes and environment, it may be a better idea to dig deeper into the pathway mechanism behind the disease. For example, some investigators have validated potential loci for association through G×E regulating exposure factors for asthma risk [1], as well as tapping into predisposing causes of difficult-to-treat diseases, including cancer [2,3], rhinitis [4], and depression [5].

However, two main methods are currently being used by breeders in agricultural production to increase crop yields and livestock productivity [6]. The first is to develop varieties with relatively low G×E effect to ensure stable production performance in different environments. The second is to use information from different environments to improve the statistical power of genomic selection (GS)and genome-wide association study (GWAS) to select individuals with better breeding value and to reveal potential loci of complex traits. The first method requires long-term commitment, while the second method clearly has faster returns. In GWAS, the use of multiple environments or phenotypes for association studies has become increasingly important. This not only improves the overall statistical power [7], but also allow to detect signaling loci for G×E. There are significant challenges when using multiple environments or phenotypes for GWAS, mainly because most diseases and quantitative traits have numerous associated loci with minimal impact [8], and thus it is impossible to determine the effect size regulated by environment in these loci. The current detection strategy for G×E is based on complex statistical model, often requiring the use of a large number of samples to detect important signals [9,10]. In GS, breeders can use whole genome marker data to identify and select target strains in the early stages of animal and plant production [11-13]. The original GS models and methods, like the GWAS models, could only analyze a single environment or phenotype [14]. To improve the predictive accuracy of the models, higher marker densities are often required, allowing the proportion of genetic variation explained by these markers to be increased, indirectly obtaining higher predictive accuracy. It is worth mentioning that the consideration of G×E and multiple phenotypes in GS models [15] has been widely studied in different plant and animal breeding [16]. GS models that allow G×E have been developed [17] and most of them have modeled and interpreted G×E using structured covariates [18]. In these studies, most of the GS models showed stronger predictive accuracy when combined with G×E compared to single environment (or phenotype) analysis. Therefore, it is necessary to develop a model that utilizes G×E information to study GWAS and GS.

This study constructed a new genotype design model program (GbyE) for G×E, and enhanced the statistical power of GWAS and GS by utilizing the interaction effects of multiple environments or phenotypes. Demonstration data from 282 maize were used for simulation experiments and real data from 354 yaks and 255 peaches were used for validation experiments to evaluate the performance of the model. The method can be embedded in any GWAS and GS analysis without discrimination.

## 2. Materials and Methods

### 2.1. Support packages

The GbyE model in this study is used in GWAS and GS, using the genome association and prediction integrated tool (GAPIT) [19], Bayesian Generalized Linear Regression (BGLR) [20], and Ridge Regression Best Linear Unbiased Prediction (rrBLUP) [21]package as support packages. In order to simplify the operation of the GbyE function package, the basic calculation package is attached to this package to support the operation of GbyE, including four function packages G&E_Simulation.R (GAPIT built-in simulation phenotype function package, with some modifications to meet the requirements of this article), GbyE.file.Calculate.R (For numerical genotype and phenotype data, this package can be used to process interactive genotype files of GbyE), Power.FDR.Calcaulate.R (Calculate the statistical power and false discovery rate (FDR) of GWAS), and Comparison.GWAS.Result.Pvalue.R (GbyE generates redundant calculations in GWAS calculations, and SNP effect loci with minimal *p*-values can be filtered by this package).

### 2.2. Samples and sequencing data

In this study, a demo data is used for software simulation analysis, and the real data of yak and peach species are used to verify the model. The demo data are from wheat of 282 inbred lines, including 4 phenotype data, and there is no missing phenotype in this data in any case. Among them, our phenotype data is simulated by using the self-made R language simulation function, and the mean value and GbyE type phenotype file are calculated. The real data come from a Jia C [22] study on growth phenotypes at weaning in yak, included genotype data of 354 yaks and 4 phenotype traits related to body weight (BW), withers height (WH), body length (BL) and chest girth (CG) at weaning, it cntains 98688 SNPs after filtering. Since the original file is in the format of ped and map, in order to facilitate subsequent analysis, plink v1.09 and self-made script are used to convert the format into HapMap format. In addition, peach data from 19 provinces, cities, and autonomous regions in China, as well as the United States, Italy, New Zealand, and Japan were added for model validation. The specific descriptive statistical data are shown in Table 2. The above data can be obtained through the methods provided in Table 1.

**Table 1.**
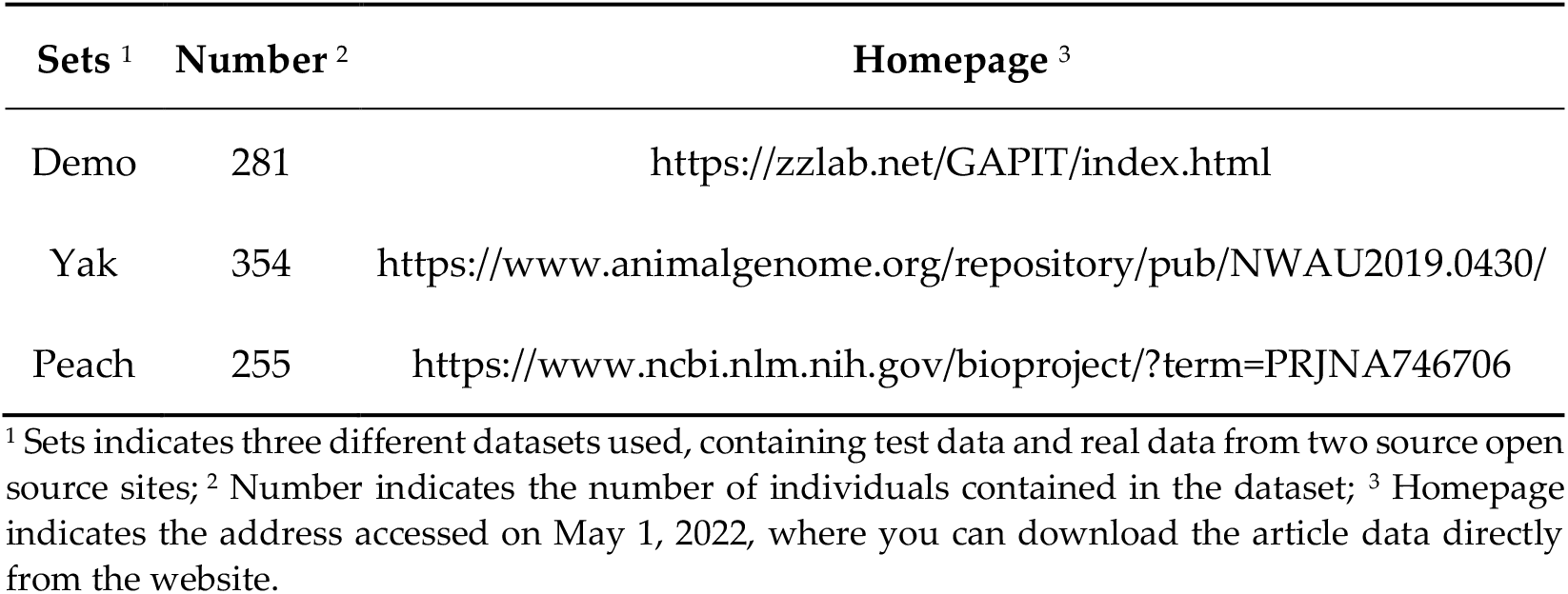
Home page of test data.

| Sets <sup>1</sup> | Number <sup>2</sup> | Homepage <sup>3</sup> |
| --- | --- | --- |
| Demo | 281 | https://zzlab.net/GAPIT/index.html |
| Yak | 354 | https://www.animalgenome.org/repository/pub/NWAU2019.0430/ |
| Peach | 255 | https://www.ncbi.nlm.nih.gov/bioproject/?term=PRJNA746706 |

<sup>1</sup> Sets indicates three different datasets used, containing test data and real data from two source open source sites; <sup>2</sup> Number indicates the number of individuals contained in the dataset; <sup>3</sup> Homepage indicates the address accessed on May 1, 2022, where you can download the article data directly from the website.

**Table 2.** Description Statistics.

| Species | Index | Phenotypes | Size | Mean | SD | Max | Min | CV |
| --- | --- | --- | --- | --- | --- | --- | --- | --- |
| Peach | P1 | Trait1 | 159 | $9.39 \times 10^{-18}$ | 1.00 | 2.16 | -1.85 | $1.07 \times 10^{17}$ |
| | P2 | Trait2 | 164 | $-1.64 \times 10^{-16}$ | 1.00 | 2.20 | -1.59 | $-6.11 \times 10^{15}$ |
| | P3 | Trait3 | 157 | $-1.82 \times 10^{-16}$ | 1.00 | 2.29 | -1.52 | $-5.50 \times 10^{15}$ |
| | P4 | Trait4 | 143 | $1.32 \times 10^1$ | 2.52 | 23.12 | 0.93 | $1.92 \times 10^{-1}$ |
| | P5 | Trait 5 | 153 | $1.95 \times 10^{-16}$ | 1.00 | 2.31 | -1.53 | $5.14 \times 10^{15}$ |
| | P6 | Trait6 | 152 | $5.69 \times 10^{-17}$ | 1.00 | 3.04 | -0.99 | $1.76 \times 10^{16}$ |
| Yak | Y1 | body weight (BW) | 350 | $8.42 \times 10^1$ | 10.30 | 117.00 | 58.00 | $1.22 \times 10^{-1}$ |
| | Y2 | withers height (WH) | 354 | $9.44 \times 10^1$ | 5.26 | 108.00 | 82.00 | $5.57 \times 10^{-2}$ |
| | Y3 | body length (BH) | 354 | $9.19 \times 10^1$ | 7.38 | 116.00 | 73.00 | $8.03 \times 10^{-2}$ |
| | Y4 | chest girth (CG) | 354 | $1.24 \times 10^2$ | 7.81 | 144.00 | 100.00 | $6.30 \times 10^{-2}$ |

\* Index indicates the index code of this phenotype in the text; number indicates the number of observations in the whole statistical phenotype file.

### 2.3. Simulated traits

Phenotype simulation was performed by modifying the G&E_Simulation simulation function of GAPIT. The function is obtained by calculating the effect value of each SNP and then randomly selecting and fitting the SNP effect from a multivariate Gaussian distribution of random numbers, so the genotype file of type 012 needs to be input during the simulation. The simulated phenotype conditions in this paper are set as follows: 1) The three levels of 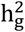are set at 0.8, 0.5, and 0.2, representing high 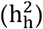, medium 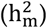and low 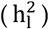heritability; 2) Genetic correlation set three levels 0.8, 0.5, 0.2 distribution representing high (R_h_), medium (R_m_) and low (R_I_) genetic correlation; 3) 20 pre-set effect loci of QTL.

### 2.4. Genetic by environment interaction model

This paper simulates the effect of genetic by environment interaction by constructing genotype files of two environments. Suppose our genotype file G is an n × m matrix, by constructing a 2 × 2, and find their Kronecker product to obtain a 2n × 2m matrix of GbyE, in which 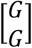matrix is defined as additive effect and 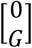matrix is defined as interaction effect. 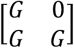matrix is called gene environment interaction matrix model. GWAS and GS analysis were performed using the original G genotype file and the constructed GbyE file.

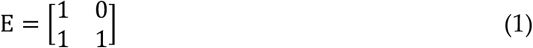

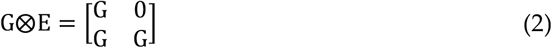

where G is the matrix of genotype files and E is the design matrix for exploring reciprocal effects. GbyE mainly uses the Kronecker product of the genetic matrix (G) and the environmental matrix (E) as the genotype for subsequent GWAS as a way to distinguish between additive and reciprocal genetic effects.

### 2.5. Association analysis model

The mixed linear model of GAPIT is used as the basic model for GWAS analysis, and the PCA parameter is set to 3. Then the *p*-values of detection results are sorted and their power and PDR values are calculated.

General expression of mixed linear model:

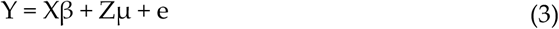

Where y is the observation vector; β is the fixed effect vector; μ Is the random effect vector, which follows the normal distribution μ∼N(0, G) with mean vector of 0 and variance covariance matrix of G; X is the incidence matrix of fixed effect; Z is the incidence matrix of random effects; e is a random error vector, and its elements need not be independent and identically distributed, e∼N(0,R). At the same time, it is assumed that cov(G, R)=0, there is no correlation between G and R, and the variance covariance matrix of y becomes:

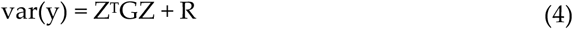

### 2.6. Power and FDR

This study uses Power and FDR indicators as evaluation items for the detection power of GbyE strategy in GWAS. And use the mean of 5-fold repeated cross validation as the reference value for statistical power comparison, compare the calculation results of GbyE with traditional Mean value calculation, additive effects, and interaction effects to evaluate the detection power of GbyE strategy. Through the comprehensive analysis of these evaluation indicators, we aim to comprehensively evaluate the statistical power of the GbyE strategy in GWAS and provide a reference for future optimization research. GbyE used the ‘Comparison.GWAS.Result.Pvalue.R’ function package to filter the calculation results of GWAS redundancy.

Among them, the formulae for calculating Power and FDR are as follows:

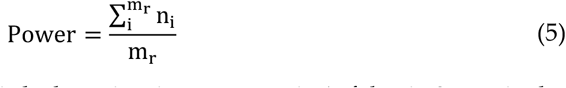

where n_i_ indicates whether the i-th detection is true, true is 1, false is 0; m_r_ is the total number of all true QTLs in the sample size; the maximum value of Power is 1.

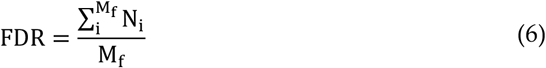

where N_i_ represents the i-th true value detected in the pseudogene, true is 1, false is and cumulative calculation; M_f_ is the number of all labeled false SNPs in the total sample; the maximum value of FDR is 1.

### 2.7. Best Linear Unbiased Prediction

Before GS, the simulated and real phenotypes were divided into test set and training set by 5-fold cross-validation, and the rrBLUP function package was used for GS, the parameter of method is “ML”. To explore the effects of additive and interactive effects in GbyE, genetic interactions were designed into different groups in BGLR. In the estimation of rrBLUP, the additive and interactive effects are regarded as a whole to estimate the effect value, which may cause weak estimates to reduce the accuracy of GS. However, the additive and interactive effects are regarded as two separate parts and estimated separately in BGLR. Relevant parameters are set as follows: 1) model set to “RRB”; 2) nIter is set to “12000”; 3) burnIn is set to “10000”. The results of the above operations are averaged over 100 cycles.

Ridge regression best linear unbiased prediction model:

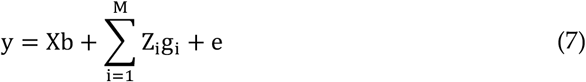

Where y is the phenotype vector; X is the fixed effect coefficient matrix, the length of which is the number of individuals in the training group, and the element value is a vector of 1; b is the fixed effect, that is, the mean phenotype of the training group; Z_i_ is the digitization of the i-th site Genotype vector, and the sum is the marker-encoded correlation matrix; g_i_ is the molecular marker effect vector of the ith locus. The effect value e is the residual error, which obeys the distribution 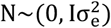.

Prediction using “RRB” model in BGLR:

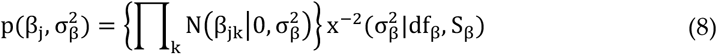

### 2.8. Bayesian Generalized Linear Regression

GS methods based on BLUP theory assume that all markers have the same genetic variance, while in reality only a few SNPs have effects genome-wide and are linked to QTLs affecting traits, and most SNPs are non-effective. When we assume the variance of marker effects as some prior distribution, the model becomes a Bayesian approach. In this paper, six Bayesian methods in BGLR were used to estimate GbyE, including BRR, BayesA, BayesB, BayesCpi, and Bayesian LASSO. In the prediction estimation of GbyE using BGLR, the numerical genotypes used in GbyE were divided into two different random effects of additive and reciprocal effects into the model, and BGLR estimated the two “lists” separately.

### 2.9. Environmental missing prediction

In this study, we simulated deletions in single and double environments using data from 281 inbred maize, 354 yaks, and 255 peach families. In the missing single environment case of the simulated experiment, we were missing phenotypes and genotypes in only one environment (GbyE had twice the number of markers by expanding genotypes to include genotypes in environment 1 and environment 2), which allowed the model to use genotype effects from environment 2 to predict phenotype values from environment 1. However, in the case of missing double environment, both phenotypes and genotypes of environment 1 and environment 2 are missing, and the model can only predict phenotypic values in the two missing environments through the effects of other markers. In addition, the data were standardized and unstandardized to assess whether standardization had an effect on the estimation of the model. This experiment was tested using the “ML” method in rrBLUP to ensure the efficiency of the model.

## 3. Results

### 3.1. GWAS statistical power of models at different heritabilities and genetic correlations

Power-FDR plots were used to demonstrate the detection efficiency of GbyE at three genetic correlation and three genetic power levels, with a total of nine different scenarios simulated (from left to right for high and low genetic correlation and from top to bottom for high and low genetic power). In order to distinguish whether the effect of improving the detection ability of genome-wide association analysis in GbyE is an additive effect or an effect of environmental interactions, we plotted their Power-FDR curves separately and added the traditional Mean method for comparative analysis. As shown in Figure 1, GbyE model can detect more statistically significant genetic loci with lower FDR under all heritability and genetic correlation combinations, followed by interaction effect, additive effect and Mean value method. However, in the combination with low heritability (Figure 1A, B, C), the interaction effect detected more real loci than GbyE under low FDR, but with the continued increase of FDR, GbyE detected more real loci than other groups. Under the combination with high heritability, all groups have high statistical power at low FDR, but with the increase of FDR, the statistical effect of GbyE gradually highlights. From the analysis of heritability combinations at all levels, the effect of heritability on interaction effect is not obvious, but GbyE always maintains the highest statistical power. The average detection power of GWAS in GbyE can be increased by about 20%, and with the decrease of genetic correlation, the effect of GbyE gradually highlights, indicating that the G×E plays a role.

**Figure 1.**
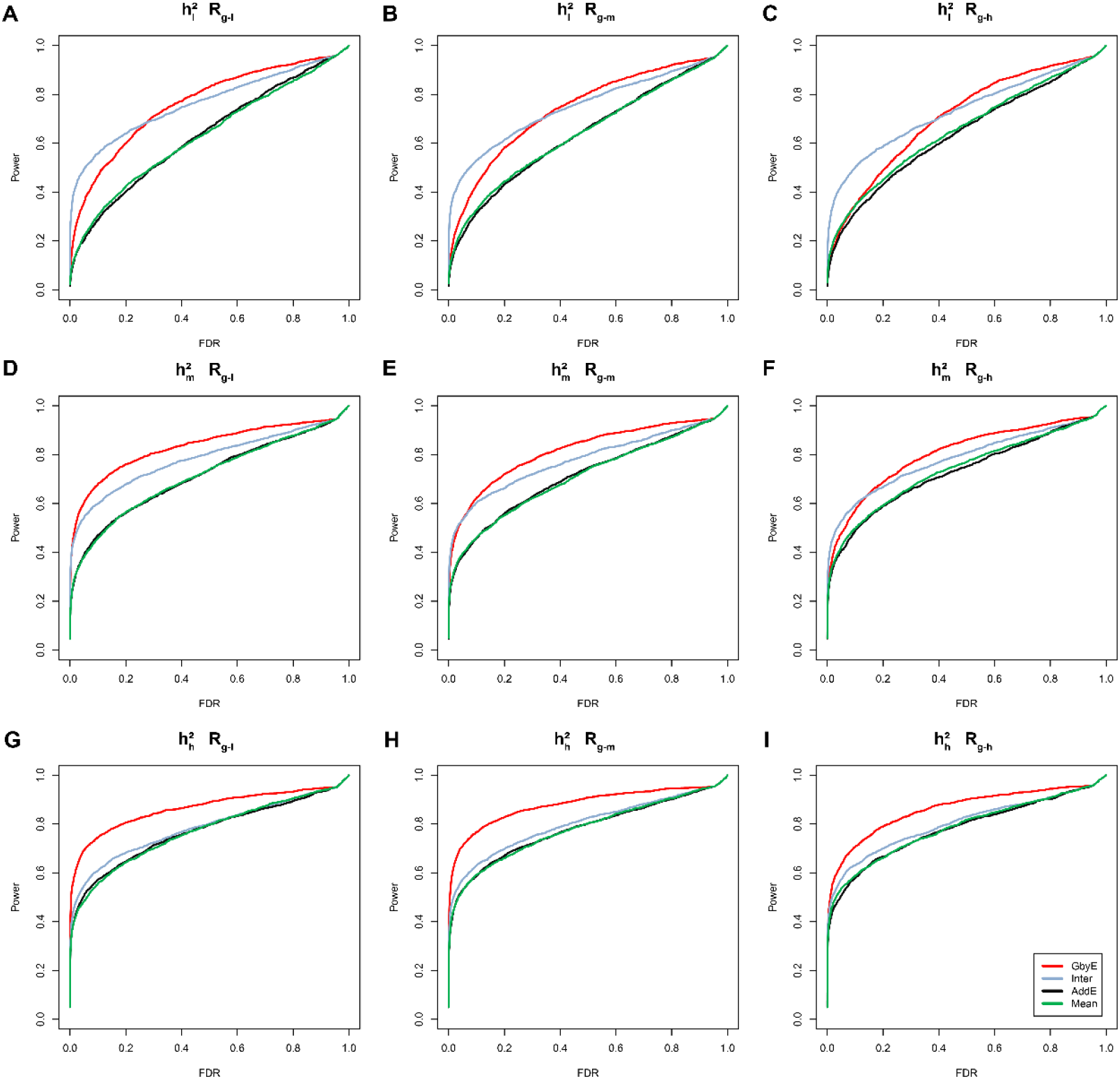
Power-FDR plot. Comparing the efficacy of the GbyE model with the conventional mean method in terms of detection power and FDR. From left to right, the three levels of genetic correlation are indicated in order of low, medium and high. From top to bottom, the three levels of heritability, low, medium and high, are indicated in order. (1) Inter: Interaction effects disentangled from GbyE; (2) AddE: Additive effects disentangled from GbyE; (3) 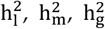: Low, medium, high heritability; (4) R_I_, R_m_, R_I_: where R stands for genetic correlation, represents three levels of low, medium and high.

### 3.2. GWAS loci detection

As shown in Figure 2, true marker loci were detected on chromosomes 1, 6 and 9 in Mean, and the same loci were detected on chromosomes 1 and 6 for the additive effect in GbyE (loci detected jointly in the two sections connected by solid lines in the figure). Although the additive effect in GbyE was not detected at loci on chromosome 9 that did not cross the significance threshold (*p*-value < 3.23×10^−6^), it has shown high significance relative to other markers of the same chromosome. In the reciprocal effect of GbyE, we detected more significant loci on chromosomes 1, 2, 3 and 10, and these loci were not detected in either of the two previous sections. The QQ plots from Figure 3 show that the overall statistical power of the additive effects in Mean and GbyE are close, but the interaction effects in GbyE reflect a higher statistical power. Overall, the model of GbyE has higher statistical power relative to the Mean and can detect more relevant loci.

**Figure 2.**
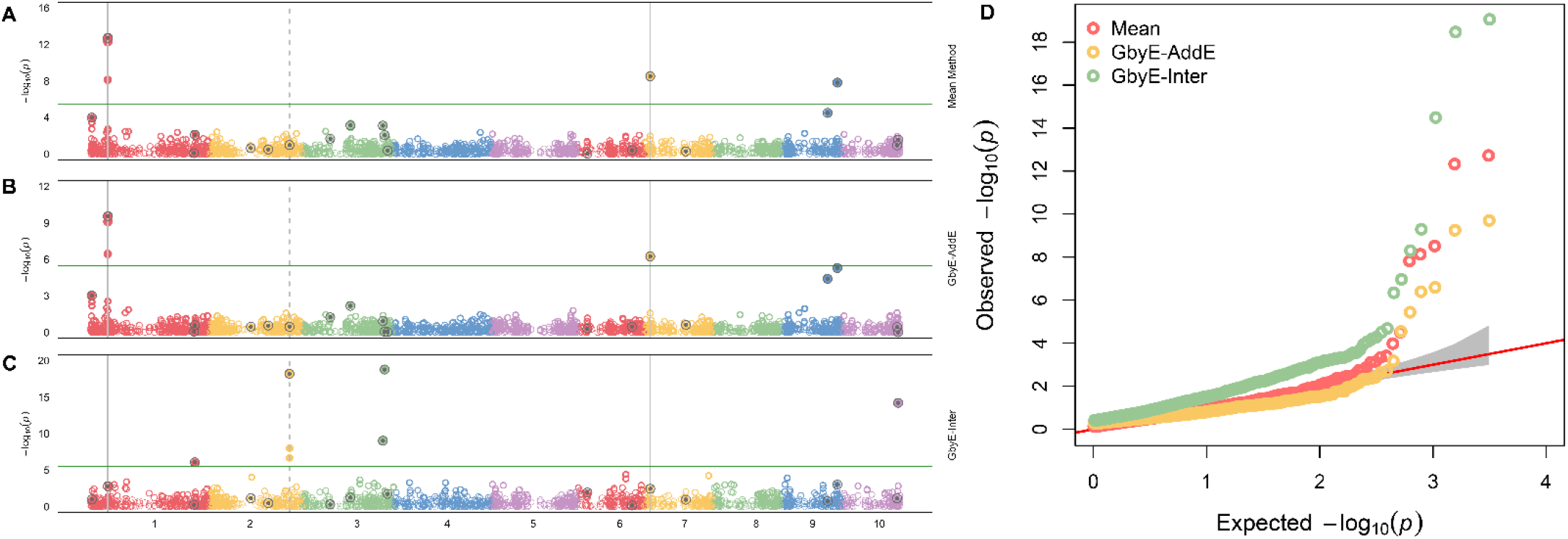
Manhattan statistical power comparison plot. Manhattan comparison plots of mean (A), additive (B) and gene-environment interaction effects (C) at a heritability of 0.9 and heritability correlation of 0.1. Different colors are used in the diagram to distinguish between different chromosomes (X-axis). Loci with reinforcing circles and centroids are set up as real QTN loci. Consecutive loci found in both parts are shown as id lines, and loci found separately in the reciprocal effect only are shown as dashed lines. Parallel horizontal lines indicate significance thresholds (*p*-value < 3.23×10^−6^). (D) Quantile-quantile plots of simulated phenotypes for demo data from genome-wide association studies. x-axis indicates expected values of log *p*-values and y-axis is observed values of log *p*-values. The diagonal coefficients in red are 1. GbyE-inter is the interaction effect in GbyE; GbyE-AddE is the additive effect in GbyE.

**Figure 3.**
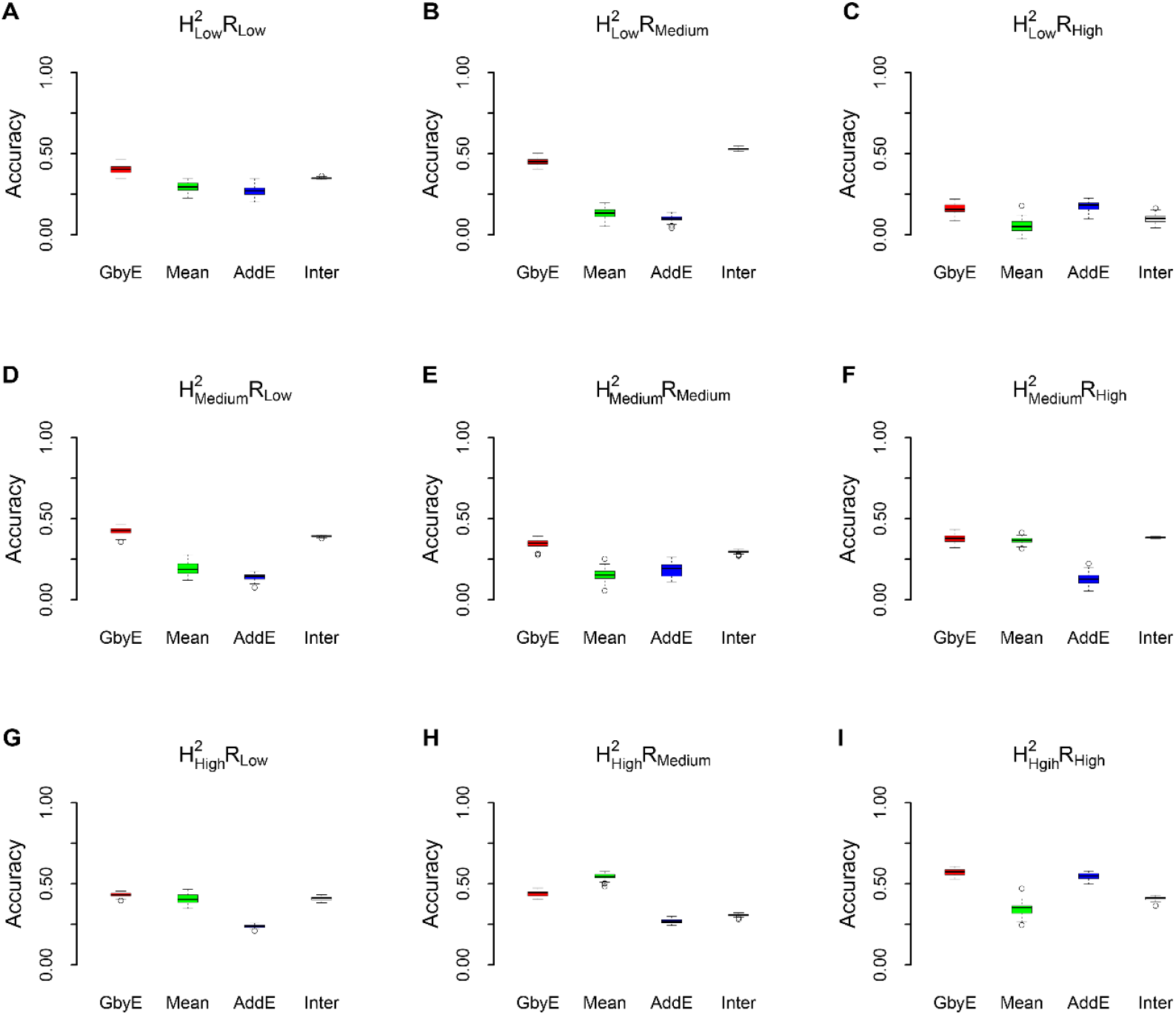
Box-plot of model prediction accuracy. The prediction accuracy (pearson’s correlation coefficient) of the GbyE model was compared with the tradition al Mean value method in a simulation experiment of genomic selection under the rrBLUP operating environment. The effect of different levels of heritability and genetic correlation on the prediction accuracy of genomic selection was simulated in this experiment. Each row from top to bottom represents low heritability 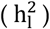, medium heritability 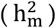and high heritability 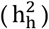, respectively; each column from left to right represents low genetic correlation (R_I_), medium genetic correlation (R_m_) and high genetic correlation (R_h_), respectively; The X-axis shows the different test methods and effects, and the Y-axis shows the prediction accuracy.

**Figure 4.**
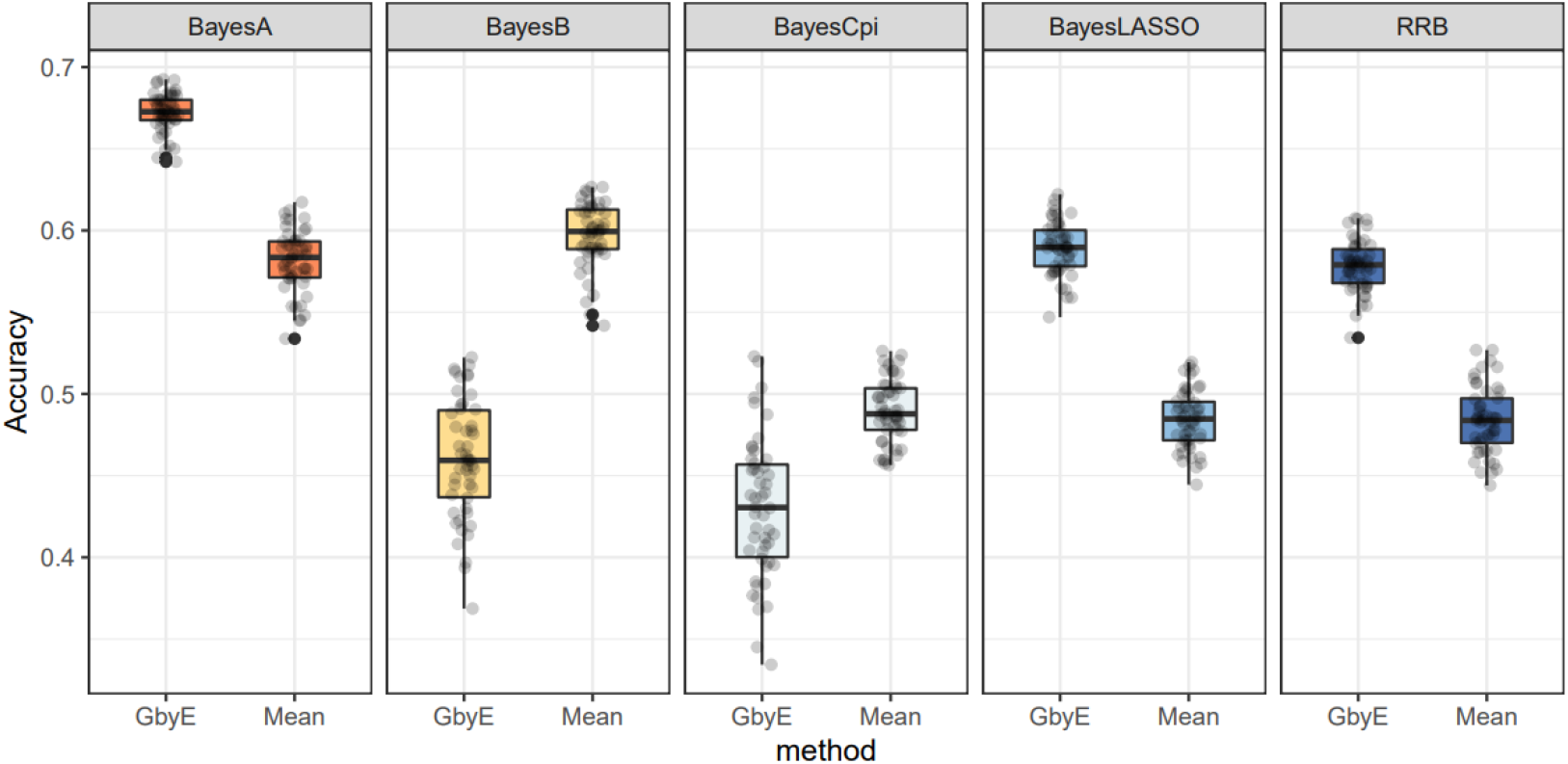
Relative prediction accuracy histogram for different Bayesian models. The X-axis is the Bayesian approach based on BGLR, and the Y-axis is the relative prediction accuracy. Where we normalize the prediction accuracy of Mean (the prediction accuracy is all adjusted to 1); the prediction accuracy of GbyE is the increase or decrease value relative to Mean in the same group of models.

### 3.3 Genomic selection of ridge regression best linear unbiased prediction

The statistical power of GbyE was significantly higher than the Mean value method by model statistics of rrBLUP in most cases of heritability and genetic correlation (Figure 3). The prediction accuracy of the additive effect was close to that of Mean value method, which was consistent with the hereditary law. The prediction accuracy of interaction effects in GbyE remains at the same level as in GbyE, and interaction effects play a role in the model. We observed that in 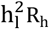(Figure 3C), 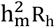 (Figure 3F), 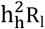 (Figure 3G), the prediction accuracy of GbyE was slightly higher than the Mean value method, but there was no significant difference overall. In addition, it was observed that the prediction accuracy of GbyE in 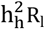(Figure 3H) was slightly lower than that of the Mean value method, with no significant difference. Under the combination of low heritability and genetic correlation, the prediction accuracy of Mean value method and additive effect model remained at a similar level. However, with the continuous increase of heritability and genetic correlation, the difference in prediction accuracy between the two gradually increases. In summary, the GbyE model can improve the accuracy of GS by capturing information on multiple environment or trait effects under the rrBLUP model.

### 3.4. Accuracy of Bayesian ridge regression

The overall performance of GbyE under the ‘BRR’ statistical model based on the BGLR package remained consistent with rrBLUP, maintaining high predictive accuracy in most cases of heritability and genetic relatedness (Figure S1). However, when the heritability is set to low and medium, the difference between the prediction accuracy of GbyE model and Mean value method gradually decreases with the continuous increase of genetic correlation, and there is no statistically significant difference between the two. The prediction accuracy of the model by GbyE in 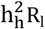(Figure S1G) and 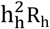 (Figure S1I) is significantly higher than that by Mean value method when the heritability is set to be high. On the contrary, when the genetic correlation is set to medium, there is no significant difference between GbyE and Mean value method in improving the prediction accuracy of the model, and the overall mean of GbyE is lower than Mean.

### 3.5. Adaptability of Bayesian models

The two methods of GbyE model and Mean value method are combined with five Bayesian algorithms in BGLR for GS analysis, and the written R script is used for phenotypic simulation test, where heritability and genetic correlation are both set to 0.5. The results indicate that among the three Bayesian models of RRB, BayesA, and BayesLASSO, the predictive accuracy of GbyE is significantly higher than that of Mean value method. In contrast, under the Bayesian models of BayesB and BayesCpi, the prediction accuracy of GbyE is lower than that of the Mean value method. The GbyE model improves the prediction accuracy of the three Bayesian models BRR, BayesA, and BayesLASSO using information from GEI and increases the prediction accuracy by 9.4%, 9.1%, and 11%, respectively, relative to the Mean value method. However, the predictive accuracy of the BayesB model decreased by 11.3%, while the BayescCpi model decreased by 6%.

### 3.6. Impact of environmental missing on the model

We tested the environmental deficiency using simulated data, 354 yak data, and 235 peaches data. And the environmental correlations between them were derived (Figure S2-3). Most of the phenotypes of the simulated data had high correlations between them, and in the yak and peach data, both environments with high and low correlations were more uniformly present, which can represent the prediction of the model in different environmental correlation cases. In the case of the demo data, we simulated a total of nine scenarios with different heritabilities and genetic correlations (Figure 5A). After analyzing the genetic correlation and heritability levels of the simulated phenotypes, no direct correlation was found between the correlation between the two environments or phenotypes used in the GbyE model and the prediction accuracy of GS, regardless of the combination of heritability and genetic correlation. Therefore, when using GbyE and Mean value methods for genomic prediction, additional factors may need to be considered to improve the prediction accuracy. The improvement in prediction accuracy by the GbyE model was found to be significantly higher than the Mean value method in single environment deletion, regardless of the combination of heritability and genetic correlation. In addition, when GbyE estimated the phenotypic values of environment 1 and environment 2 alone, its prediction accuracy was too high and the overfitting phenomenon that existed. This means that GbyE is affected by noise or randomness in the training data, which leads to over-fitting to a specific environment. GbyE does not show a significant decrease in GS prediction accuracy improvement when both environments are missing. On the contrary, the prediction accuracy significantly decreases when GbyE was estimated for environment 1 and environment 2 alone (Figure 5B).

**Figure 5.**
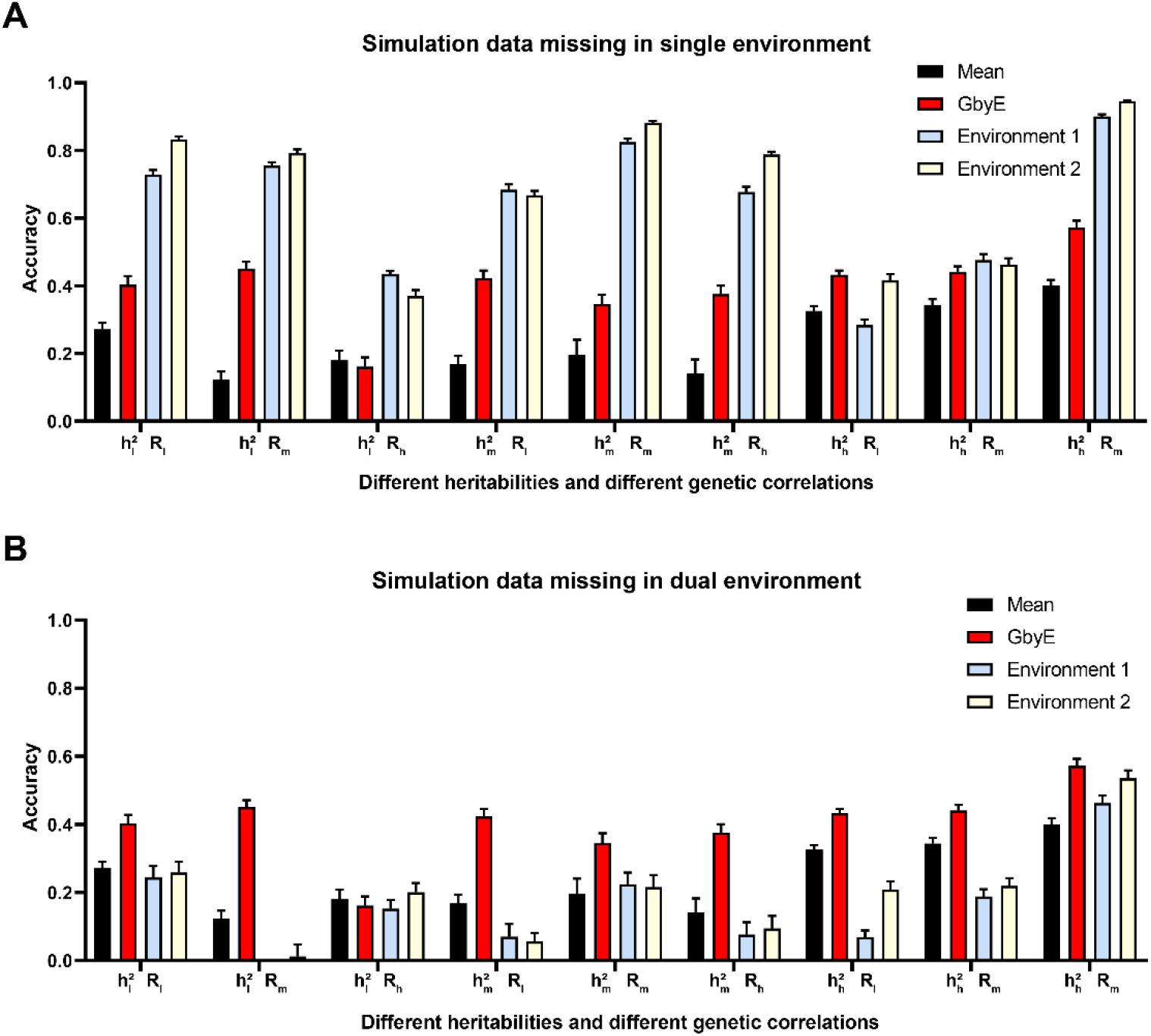
Prediction accuracy of simulated data in single and dual environment absence. The prediction effect of GbyE was divided into two parts, environment 1 and environment 2, to compare the prediction accuracy of GbyE when predicting these two parts separately. This includes simulations with missing phenotypes and genotypes in environment 1 only (A), and simulations with missing in both environments (B). The horizontal coordinates of the graph indicate the different combinations of heritabilities and genetic correlations of the simulations.

Whether the phenotypic data of yak and peach were standardized or not may significantly affect the prediction results of GS. For the peach data with a single environment missing and treated with phenotypic normalization (Figure 6A), it was found that when there is a high correlation between Environment 1 and Environment 2, GbyE predicts Environment 1 or Environment 2 alone with higher prediction than the Mean value method, but the overall prediction accuracy of GbyE is lower. As the environmental correlation gradually decreases, the predictive accuracy of the model also gradually decreases. When the above test groups are not standardized (Figure 6B), most results are consistent with the former, but the GbyE model shows overfitting under low correlation. In addition, when both environments were missing (Figure 6C, D), mean value method was signififcantly higher than GbyE that predicts environment 1 or environment 2 alone, which is consistent with the simulated data results. For the analysis of yak data, when a single environment was missing and the data were normalized (Figure 6E), although the overall prediction accuracy of GbyE was not significantly improved, in most environmental combinations, the accuracy of Environment 1 and Environment 2 alone was significantly higher than the Mean value method. Prediction accuracy when predicting environments 1 and 2 alone did not differ significantly between those without and with standardization (Figure 6F, H). However, when GbyE is not standardized, the square value of some prediction accuracy is much higher than heritability (Y1-Y2, Y1-Y3 and Y3-Y4), showing a overfitting situation. In the experiment of dual environment missing prediction (Figure 6G, H), the overall prediction results were consistent with the previous results.

**Figure 6.**
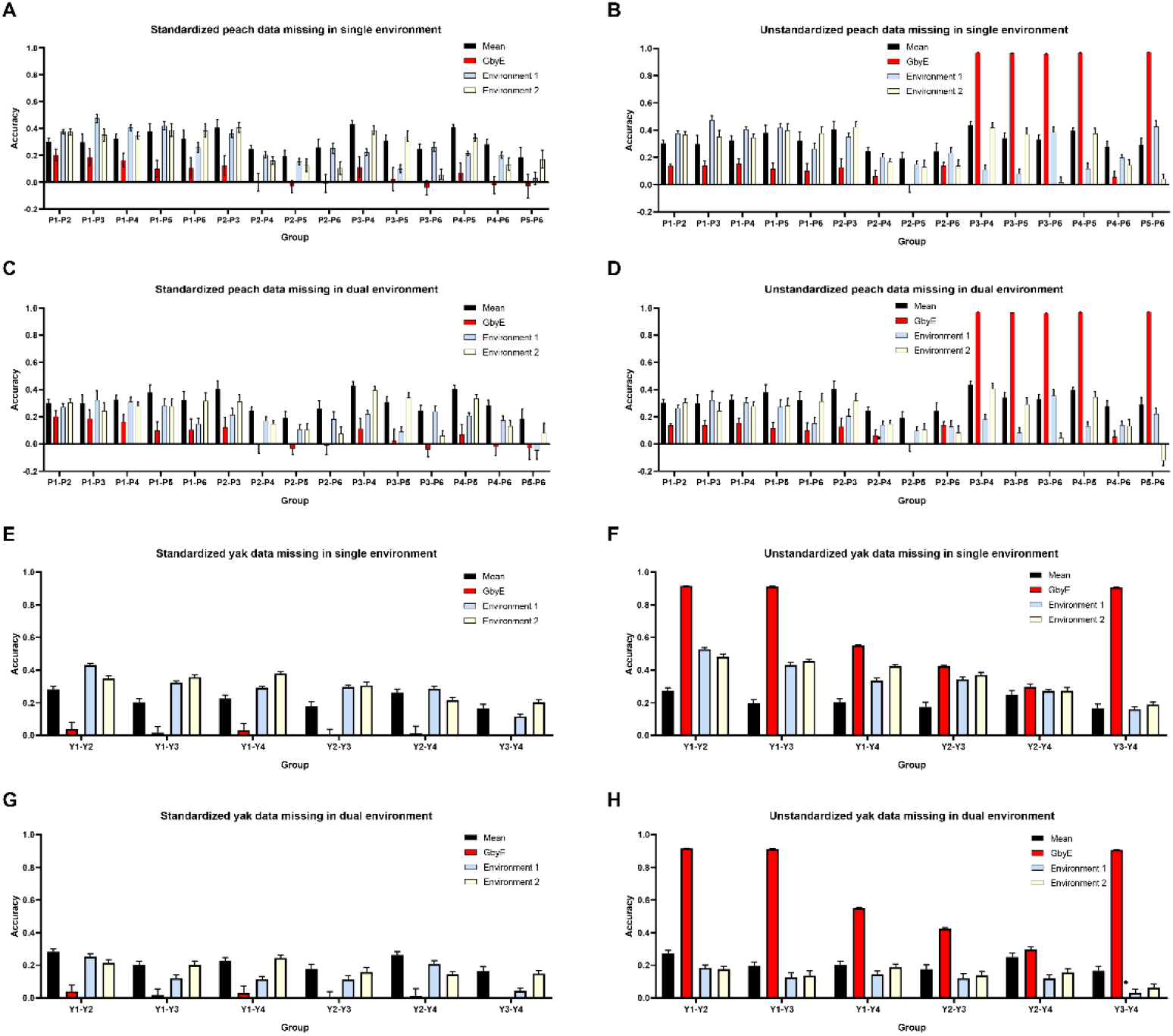
Box line plot of prediction accuracy and environmental relevance for real data. Two environments were predicted separately for peach and yak data using GbyE. Single-environment missing and dual-environment missing were considered, as well as whether the data were standardized or not. (A) standardized peach data missing in a single environment; (B) unstandardized peach data missing in a single environment; (C) standardized peach data missing in a dual environment; (D) unstandardized peach data missing in a dual environment; (D) standardized yak data missing in a single environment; (F) unstandardized yak data missing in a single environment; (G) standardized yak data are missing in both environments; (H) unstandardized yak data are missing in both environments.

## 4. Discussion

This study constructed an interaction model of GbyE that takes into account genetic and environment interactions and multi-trait information and can help to improve the statistical power of GWAS and GS. The phenotype of organisms is usually composed of multiple factors, mainly genetic [23] and environmental factors [24], especially the phenotype of quantitative traits is often influenced by a combination of these factors [25,26]. The additive effects included in GbyE are essentially equivalent to the computational process of traditional models, but the interaction effect in GbyE can enable GWAS to detect more real loci with a lower FDR. In addition, the prediction accuracy of GS is often affected by genetics, environmental variation [27], polygenic effects [28], sample size and quality [29], and model selection [30]. Currently in GS, the accuracy of prediction will decrease if there are relatively large genetic background differences in the test population from the training data. Since genomic predictions can only predict the effects of genetic and fixed environment, but not the random residuals, the square of the prediction accuracy cannot exceed the level of heritability, which is also called heritability limitation. Detecting two genetic effects (additive and interactive effects) in GWAS and GS is a boost to computational complexity, while obtaining genotypes for genetic interactions by Kronecker product is an efficient means. This allows the estimation of additive and interactive genetic effects separately during the analysis, and ultimately the estimated genetic effects for each GbyE genotype (including additive and interactive genetic effect markers) are placed in a t-distribution for *p*-value calculation, and the significance of each genotype is considered by multiple testing.

The genetic correlation in the simulated phenotype function defines different levels. When the genetic correlation level is set high, then additive genetic effects will dominate, and interactive genetic effects with different traits or environments are often at lower levels [31]. Therefore, the statistical power of the GbyE model did not improve significantly compared with the traditional Mean value method when simulating high levels of genetic correlation. On the contrary, in the case of low levels of genetic correlation, the genetic variance of additive effects is relatively low and the genetic variance of interaction effects is dominant. At this time, GbyE utilizes multiple environments or traits to highlight the statistical power. Since the GbyE model obtains G×E information by encoding numerical genotypes, it only increases the volume of SNP data and can be applied to any traditional GWAS association statistical model. However, this may slightly increase the correlation operation time of the GWAS model, but compared to other multi environment or trait models [32,33], GbyE only needs to perform a complete traditional GWAS once to obtain the results. Traditional models are more complex, lengthy, and prone to errors when making makers selection, which can affect the results of association prediction. And we require multi environment or trait models to have a simpler computational structure than single trait models to overcome the uncertainty associated during estimation.

In GS, rrBLUP algorithm is a linear mixed model-based prediction method that assumes equal effects for all genes and no pleiotropy [34]. This algorithm estimates the predicted values by solving system of linear equations, assuming that the effects of all genes come from a common normal distribution with a specific value of variance, which is relatively fast to compute, but ignores the variability among genes. In contrast, the BGLR model is a linear mixed model based on Bayesian methods, which assumes that gene effects are randomly drawn from a multivariate normal distribution and genotype effects are randomly drawn from a multivariate Gaussian process, which takes into account potential pleiotropy and polygenic effects and allows inferring the effects of single gene while estimating genomic values [28]. The algorithm typically uses Markov Chain Monte Carlo methods for estimation of model parameters [35,36]. The model has been able to take into account more biological features and complexity, and therefore the overall improvement of the GbyE model over Mean is smaller under BGLR. In addition, the length of the Markov chain set on the BGLR package is often above 20,000 to obtain stable parameters and to undergo longer iterations to make the chain stable [37]. GbyE is effective in improving the statistical power of the model under most Bayesian statistical models. In the case of the phenotypes we simulated, more iterations cannot be provided for the BayesB and BayesCpi models because of the limitation of computation time, which causes low prediction accuracy. It is worth noting that the prediction accuracy of BayesCpi may also be influenced by the number of QTLs [38], and the prediction accuracy of BayesB is often related to the distribution of different allele frequencies (from rare to common variants) at random loci [39].

The overall statistical power of GbyE was significantly higher in missing single environment than in missing double environment, because in the case of missing single environment, GbyE can take full advantage of the information from the second environment in that individual. And the correlation between two environments can also affect the power of the GbyE model in different ways. On the one hand, a high correlation between two environments can improve the predictive accuracy of the GbyE model by using the information from one environment to predict the genetic values of the genome in the other environment, even if there is no direct data about that environment [40,41]. On the other hand, when two environments are extremely uncorrelated, GbyE model trained in one environment may not export well to another environment, which may lead to a decrease in prediction accuracy [42]. In tests with real data, the overall predictions for yaks appear to be overfitted, possibly because the GbyE model uses a GS model trained in one environment and tested in another environment, and the high correlation between environments may resulit to the model capturing similarities between environments unrelated to G×E information [43]. However, when estimating the breeding values for each environment separately, GbyE still made effective predictions using the genotypes in that environment and maintained high prediction accuracy. As expected, environment 2 has been designed as a marker with additive effects, and its predictive accuracy has always been consistent with the Mean value method. environment 1, as a design marker for interaction effects, has a stronger ability to perceive the environment, and changes in environmental factors have a direct impact on the predicted results. Therefore, the impact of environment 1 on prediction accuracy is more significant when two environments are missing. The correlation between the two environments may have both positive and negative effects on the power of the GbyE, so it is important to carefully consider the relationship between the two environments in subsequent in development and testing.

A key advantage of the GbyE model is that it can be applied to almost all current genome-wide association and prediction models, meaning that GbyE can significantly improve the accuracy of genomic predictions in most cases. The GbyE model may have implications for the design of future GS studies. For example, the model could be used to identify the best environments or traits to include in GS studies in order to maximize prediction accuracy. It is particularly important to test the model on large datasets with different genetic backgrounds and environmental conditions to ensure that it can accurately predict genome-wide effects in a variety of contexts.

## 5. Conclusions

GbyE simulates the effects of gene-environment interactions by building genotype files for multiple environments or multiple traits, normalizing the effects of multiple environments and multiple traits on marker effects, by using one model for genome-wide association and prediction. GbyE enables higher statistical power for GWAS and GS in any genetic power and genetic correlation. The AddE part of GbyE maintains the same statistical power as traditional Mean value method with the same statistical power and is able to detect more real loci. In addition, GbyE can adapt to most Bayesian models, increasing the statistical power of three Bayesian models, BRR, BayesA, and BayesLASSO. GbyE can make full use of multiple environment information to enhance the statistical power of the model, which can help us understand the interaction between genes and environment and provide a method for predicting association loci of complex traits.

